# Mapping Cell Fate Transition in Space and Time

**DOI:** 10.1101/2024.02.12.579941

**Authors:** Yichen Gu, Jialin Liu, Chen Li, Joshua D. Welch

**Author notes:** Equal Contribution (Yichen Gu, Jialin Liu).

## Abstract

Cell fate transition is fundamentally a spatiotemporal process, but previous work has largely neglected the spatial dimension. Incorporating both space and time into models of cell fate transition would be a key step toward characterizing how interactions among neighboring cells, the presence of local niche factors, and physical migration of cells contribute to tissue development. To realize this potential, we propose topological velocity inference (TopoVelo), a model for jointly inferring spatial and temporal dynamics of cell fate transition from spatial transcriptomic data. TopoVelo extends the RNA velocity framework to model single-cell gene expression dynamics of an entire tissue with spatially coupled differential equations. Our principled probabilistic approach enables the incorporation of time point labels and multiple slices. We further introduce the idea of cell velocity, which is defined as the physical direction of cell maturation and migration. Simulated data analysis indicates that incorporating spatial coordinates significantly improves the accuracy of velocity and time inference. When evaluated on real Slide-Seq and Stereo-Seq data, TopoVelo significantly improves the spatial coherence of inferred cell ordering compared to previous methods. Furthermore, TopoVelo accurately recovers the expected directions of cell differentiation and migration in the embryonic mouse cerebral cortex, gut, and lung. Our work introduces a new dimension into the study of cell fate transitions and lays a foundation for modeling the collective dynamics of cells comprising an entire tissue.

## 1 Introduction

Cell fate transitions are fundamentally spatiotemporal processes regulated by factors that vary over space, including secreted signaling factors that diffuse to nearby cells and bind receptors, extracellular matrix interactions, and tight junctions between adjacent cells. Clonal relationships between progenitor and descendant cells are also spatial relationships: As cells divide over time, daughter cells arise in locations spatially adjacent to the mother cell. Moreover, cells often develop in defined spatial directions over time–for example, stratified epithelium forms from basal layer to luminal layer[1,2]. During early development, cells must migrate to specific locations to form unique tissues and organ systems. For instance, the layers of the cerebral cortex form as cells migrate along radial glial processes from the ventricular zone to the outside of the developing cortex[3,4,5]. Inhibitory interneurons also migrate into the cortex from the medial and caudal ganglionic eminence[6,7,8]. Recent breakthroughs in modeling human post-implantation embryo stages have been driven by the self-assembling properties of distinct lineages mixed together that signal to each other[9]. This again highlights the critical role of spatial context in organ development.

Spatial transcriptomics enables whole-transcriptome or targeted measurement of gene expression and spatial position at cellular or sub-cellular resolution ([10,11,12,13]), thereby affording a layer of context that is typically lost in techniques that require cell dissociation, such as single-cell RNA sequencing[14,10,15,16]. Other spatial transcriptomic technologies can measure selected genes at single-molecule resolution using multiplexed in situ hybridization [17,18,19]. Spatial transcriptomics can measure not only a single slice of tissue but multiple slices from the same 3D sample at a single time point or multiple samples across successive time points. This novel technology provides the data we need to address a fundamental biological question - how are time and space linked during cell fate transition? However, no existing computational approaches can directly address this question. Pseudotime methods[20,21,22] do not model spatial positions, cannot infer direction or rate of change, and are not mechanistic. Approaches based on optimal transport[23,24] can learn a spatial or temporal coupling between two groups of cells, but they require multiple time points, cannot be applied to single-slice data, and are not mechanistic. RNA velocity methods[25,26,27,28,29,30] can infer directions and rates of change by fitting interpretable rate parameters, but do not consider spatial information. Thus, we would like to develop a computational approach that is mechanistic, models spatial coupling among cells, and can incorporate time labels and multiple sections.

Here, we present Topological Velocity Inference (TopoVelo), a model for jointly inferring the dynamics of cell fate transition over time and space. We extend the RNA velocity framework to jointly model the differentiation of all cells in a tissue using spatially coupled differential equations, formulate a principled Bayesian latent variable model that describes the data generation process, and derive an approximate Bayesian estimation method using autoencoding variational Bayes. This allows us to infer spatially coupled cell differentiation stages and rate parameters of gene expression. Additionally, we can infer the physical directions and rates of cell maturation and migration–a quantity we term “cell velocity”. We verified our method on a number of both simulated and real datasets and visualized cell velocity on spatial coordinates. Our results show that modeling spatial coupling improves cell velocity inference and brings a new dimension to the modeling of cell fate transition.

## 2 Methods

### 2.1 Problem Statement

Our goal is to use spatial transcriptomic data to infer how the gene expression levels of an entire population of cells change together during cell fate transitions. That is, we are interested in the following problem. Each sample (cell or spot), indexed by *i*, is represented by a vector of gene expression values *X*_*i*_(*t*) *∈* ℝ^*d*^ and spatial coordinates (*x, y*), both parametrized by developmental stage (time) *t*. The trajectory *X*_*i*_(*t*) is governed by some differential equation plus random noise. For each *i*, only the vector *x*_*i*_ := *X*_*i*_(*t*_*i*_) is observed at some unknown time *t*_*i*_. Furthermore, the parameters of the differential equations are spatially coupled in an unknown way, forming a random field. Our goal is two-fold: recover the latent times *t*_*i*_ for each sample and predict future gene expression values and spatial positions, i.e., *X*_*i*_(*t*) for *t > t*_*i*_.

This problem arises from the limitations of spatial transcriptomic technologies, which require fixing, freezing, and/or sectioning the cells and thus cannot measure gene expression longitudinally on the same cells. Another aspect of this problem is cell-to-cell variation in the rate of fate progression–at a given moment, cells in a population will be at different developmental stages. Thus, latent developmental time is often more informative than clock time when studying cell fate transitions.

To approach this problem, we make a few key modeling decisions. First, we describe the system using a differential equation that reflects the biochemical steps required for gene expression: a gene is transcribed as unspliced pre-mRNA before being converted into spliced mature mRNA. This builds on previous work from our group and others[31,25,32,27]. Second, we model the random field of cells using a graph that connects cells nearby in space. The spatial transcriptomic dataset can be described as a graph in which cells or spots are nodes, gene expression levels and spatial positions are node features, and edges represent spatial proximity relationships. More formally, let ***U*** and ***S*** be cell-by-gene count matrices of unspliced and spliced mRNA and ***L*** = *{*(*x*_*i*_, *y*_*i*_)*}* be spatial coordinates obtained from a spatial transcriptomic experiment. We describe the dynamical system underlying this count generation process using the following system of differential equations:

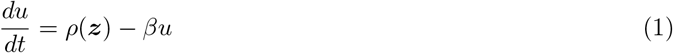

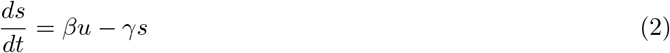

where *ρ, β* and *γ* are transcription, splicing and degradation rates. Following our previous VeloVAE paper[27], *ρ* is a transcription rate parameter that varies with cell state (***z***) and is unique to each cell and each gene, while *β* and *γ* are specific to each gene but shared across all cells. Note that *u* and *s* are the only variables in these equations that are directly observed, while cell time *t* and cell state ***z*** are latent.

We represent the population of cells as a graph *𝒢* = (*V, E*), where *V* = [*n*] is a set of nodes representing cells and *E* contains the neighboring relations between any two cells. We denote the set of neighboring cells of *i*, including *i*, by *nbr*(*i*). For each node (cell) *i*, we take its unspliced and spliced mRNA read counts as its node features, denoted as ***o***_***i***_. Importantly, we assume that this graph captures spatial dependence among the ***U***, ***S***, and *ρ* parameters of the cells. By default, we do not assume any edge features, but these could be added to the graph if available.

Using this differential equation and graph structure, we can describe how the gene expression levels vary given rate parameters, cell state, and cell time. The ODE system has an analytical solution, so if we knew the cell state and cell time values, we could readily estimate the parameters. However, cell state and cell time are latent. Thus, we can reduce the modeling problem to a latent variable inference problem. We denote ***Ƶ*** = *{****z***_***i***_ : *i ∈* [*n*]*}* and ***𝒯*** = *{t*_*i*_ : *i ∈* [*n*]*}* as the set of all cell states and time. We next show how to perform Bayesian estimation of ***Ƶ*** and ***𝒯*** using autoencoding variational Bayes. Intuitively, our TopoVelo approach extends the RNA velocity framework to spatial transcriptomic data by incorporating cell position with graph neural networks (GNN).

### 2.2 TopoVelo: Topological Velocity Inference from Spatial Transcriptomic Data

#### Probabilistic Model

To solve the above computational problem, we first formalize our probabilistic model of the data generation process. Next, we solve the inference problem. Our generative model is a probabilistic graphical model. Different from many previous graphical models[27,29,30], which consider data generation of all cells as independent and identically distributed, we consider data generation of all cells as coupled and jointly distributed. We assume cells are nodes in a graph connected by their spatial adjacency relations and node features ***𝒪*** := *{****o***_***i***_ : *i ∈* [*n*]*}* are generated from ***Ƶ*** and ***𝒯*** via a spatially-coupled dynamical system:

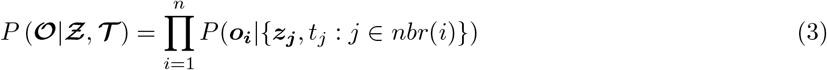

Note that the observation ***o***_***i***_ depends on its spatial neighbors *nbr*(*i*). We further assume *P* (***o***_***i***_|*{****z***_***j***_, *t*_*j*_ : *j ∈ nbr*(*i*)*}*) is a factorized Gaussian distribution over all genes, i.e.,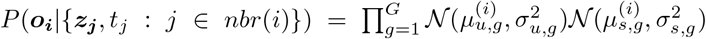 where *𝒩* (*·*) denotes a Gaussian distribution with parameters inside the parentheses. Here, we assume gene-wise variances and the variance of each gene is the same across all cells. The transcription rate for each gene *g* in each cell *i* is a function of both the cell state of *i* and the cell states of the neighboring cells in the tissue, *nbr*(*i*). The splicing and degradation rates are different for each gene, but shared across cells, giving the following system of ODEs:

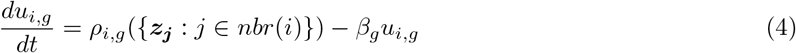

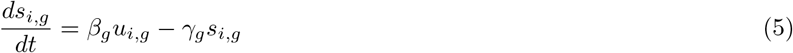

In practice, we find that it is helpful to explicitly model repression-only genes (for which *ρ* is identically 0), so we introduce an indicator variable *k*_*g*_, which is 0 if a gene is only repressed and 1 otherwise. So now the transcription rate 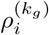 is also a function of *k*_*g*_, and in particular 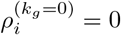for all *i*. We model each gene as a Gaussian mixture of the two states *k*_*g*_ = 0 and *k*_*g*_ = 1. We use ***w***_***g***_ to denote a one-hot encoding of *k*_*g*_ and assume it is categorically distributed: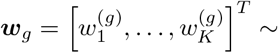 Categorical 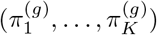, where *K* is the total number of categories and *K* = 2 in our case. We then set a Dirichlet prior,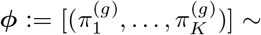 Dirichlet(*ψ*_1_, …, *ψ*_*K*_). Define ***w*** := *{****w***_***g***_ : *g ∈* [*G*]*}*. Thus Eq.3 becomes:

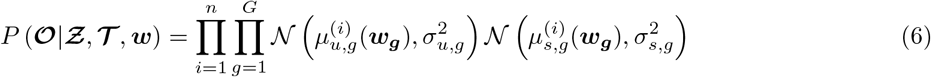

Exact inference of the latent variables ***𝒯*** and ***Ƶ*** from this model is intractable. Alternative approaches such as expectation-maximization are difficult in this setting because observations are no longer i.i.d., and we also would like to model the parameter uncertainty rather than obtaining point estimates. We thus pursue a variational Bayesian approximation. Because we want to use a graph neural network to represent a general nonlinear spatial correlation among neighboring cells, autoencoding variational Bayes [33] is a natural choice. This approach optimizes the evidence lower bound (ELBO) to jointly train an inference model (encoder network) and generative model (decoder network). To derive the ELBO in our setting, we use a variational posterior over all cell states and time, *q*(***Ƶ, 𝒯*** |***𝒪***), to approximate the true posterior. As is standard with autoencoding variational Bayes, we use a factorized Gaussian as our variational distribution, i.e.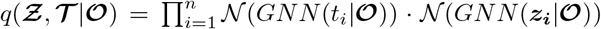. Note that the factorized distribution implies conditional independence of latent variables given the entire graph, but does not violate our spatial correlation assumption. To account for spatial correlation in the generative model, we use a GNN to map latent cell states to cell-wise transcription rates *ρ*_*i*_.

Following similar arguments to those used in previous autoencoding variational Bayes settings [33,34,35], we can derive the following evidence lower bound (ELBO) for our model:

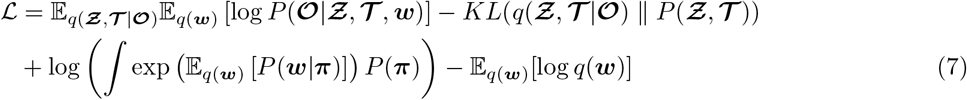

We can then perform approximate Bayesian inference by using stochastic minibatch gradient descent to optimize the ELBO with respect to the neural network parameters and ODE parameters.

#### Predicting Cell Velocity

Previous work introduced the concept of RNA velocity, defined as the time derivative of spliced mRNA,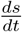. Because we know the spatial coordinates (*x, y*) of cells in our spatial transcriptomic data, we can use our modeling framework to quantify temporal changes in cell position. In particular, we can infer 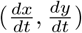, the time derivative of each cell’s spatial coordinates. We refer to the quantity 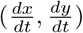simply as “cell velocity”, because it is a change of physical position over time–a velocity in the traditional sense. There are at least two possible methods for inferring cell velocity: (1) use the heuristic method popularized by scVelo to project RNA velocity onto spatial coordinates and obtain an implicit estimate of cell velocity or (2) train a graph neural network to obtain an explicit parametric form for cell velocity as a function of cell state and time. We refer to the second approach as a “spatial decoder”. One potential advantage of the spatial decoder is that it can perform out-of-sample prediction of cell positions. We compared both approaches in this paper. See below for details of spatial decoder approach.

#### Neural Network Architecture and Training

We implement TopoVelo using an encoder-decoder neural network framework. The network takes a graph as input, with nodes representing cells or spots and node features indicating spliced and unspliced counts and spatial coordinates. Edges in the graph represent spatial proximity. The encoder neural network uses graph convolutional layers to estimate the cell state and time of each cell. The decoder network uses graph convolution to estimate the transcription rate of each gene in each cell, then uses the analytical solution to the ODE to estimate the splicing and degradation rate for each gene. The node features are then reconstructed using the ODE solution given time and ODE parameters. Fig. 1b summarizes the model architecture. Our approach is agnostic to the choice of graph convolution, and in our experiments, we compare two popular graph convolution approaches, spectral graph convolution(GCN) [36] and graph attention(GAT) [37]. Both types of graph convolution compute the representation of a node using its own and its neighbors’ features.

**Fig. 1.**
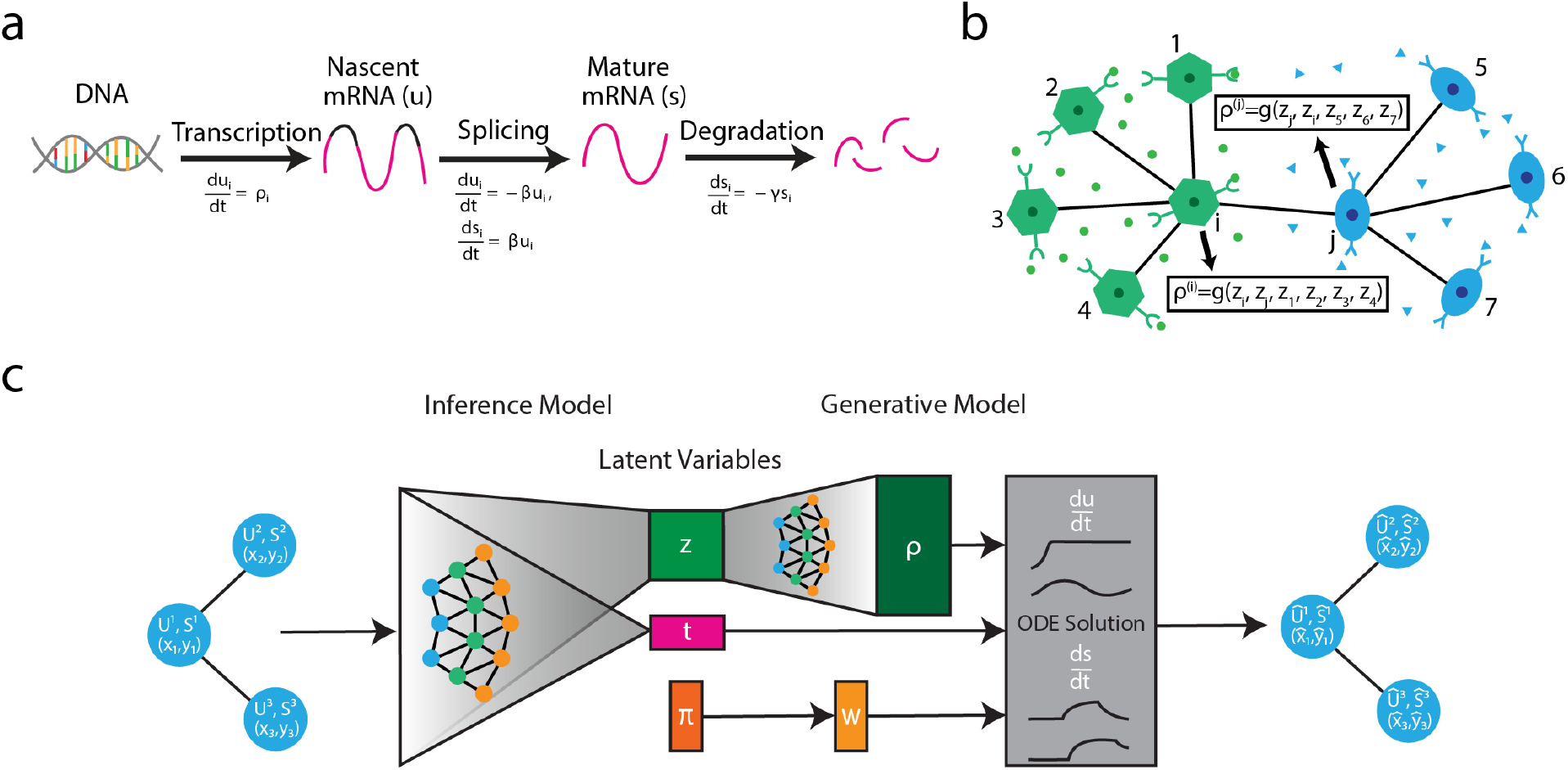
Diagram of TopoVelo model. **(a)** System of differential equations that describes how unspliced pre-mRNA (*u*) and spliced mature mRNA (*s*) values are related during gene expression. *ρ* is a transcription rate parameter that is different for every cell and every gene. *β* and *γ* are splicing and degradation rates, which are different for each gene but shared across cells. **(b)** Diagram of spatial dependence in rate parameters. TopoVelo extends previous RNA velocity models by allowing transcription rate parameters to be a function of the states of neighboring cells in a tissue, which may influence each other through mechanisms such as ligand-receptor signaling. **(c)** Diagram of TopoVelo neural network architecture. TopoVelo takes a graph as input. In this graph, nodes represent cells or spots; *u, s*, and spatial coordinates are node features; and edges represent spatial proximity. A graph neural network infers cell state and cell time jointly across cells (encoder network). Another graph neural network predicts transcription rates *ρ* from the states of each cell and its neighbors. The phase of each gene is also predicted (*w*). The *u* and *s* counts are reconstructed using the analytical solution to the differential equation from (a). The spatial positions are reconstructed using a separate graph neural network trained after the encoder is frozen.

##### Model architecture

The encoder consists of a GCN or GAT layer and a fully connected (FC) layer. It first extracts hidden features from node features ***𝒪*** and graph *G*. Next, the FC layer maps the hidden features to the mean and standard deviation of a variational posterior. The decoder is a combination of GNN and ODE solution. First, we use a decoder GNN, with a GCN or GAT layer plus an FC layer, to learn a mapping from latent cell states to transcription rates ***ρ***. Next, we compute *u* and *s* using the analytical solution to the ODEs (4)-(5), as reported in our previous work [27]. To model spatial coordinates as a function of time, we train a graph neural network after the encoder-decoder network for reconstructing *u* and *s* has converged. The spatial decoder is a graph neural network with two GCN or GAT layers and a final fully connected layer. The input to our spatial decoder is a spatial graph, with latent cell state, ***z***, latent cell time, *t*, initial time *t*_0_ and initial location (*x*_0_, *y*_0_) as node features and the output is two-dimensional spatial coordinates. (*x*_0_, *y*_0_) represent the location of a cell at a previous time, *t*_0_. We estimate *t*_0_ and (*x*_0_, *y*_0_) by taking the average time and coordinates of nearest neighbors in the cell state space from all cells within certain radius.

##### Parameter initialization

The graph neural network weights are initialized by a Xavier uniform distribution, while the weights of fully connected layers are initialized by a Xavier normal distribution. ODE parameters are initialized by the steady-state rate parameter estimates, as in our previous work [27]. To initialize *q*(***w***), we first follow scVelo and use a the dynamical model to initialize cell time. Then, we cluster genes by their co-expression and determine if they are repressed using a set of Kolmogorov-Smirnov tests. Please refer to the appendix for details.

##### Prior Distributions

By default, TopoVelo uses a multivariate Gaussian distribution with diagonal covariance matrix as a prior of latent cell state and time, *P* (***Ƶ, 𝒯***), i.e. all dimensions of cell state and cell time of all cells are mutually independent. However, when the capture times of cells are known, we can incorporate this information as an informative prior distribution for the latent cell time *t*. We assume the prior time distribution is a Gaussian with the capture times, e.g. embryonic day 10.5 or 11.5, as the mean and a variance proportional to the time intervals between adjacent experiments.

We train TopoVelo using gradient descent with the ADAM optimizer [38]. Because we reconstruct the node features of the graph but not the edges, we can enable mini-batch training to reduce memory consumption greatly. After training TopoVelo, we concatenate the latent time, cell state, initial time and initial location into feature vectors and use the given spatial coordinates as target outputs to train our spatial decoder, also using stochastic mini-batch gradient descent with the ADAM optimizer.

## 3. Results

### 3.1 Modeling Spatial Positions Improves Time and Velocity Inference from Simulated Data

We first investigated the performance of TopoVelo using simulated data and compared it with previous RNA velocity inference methods, which do not utilize spatial information. To simulate cells from an entire tissue, we extended scVelo’s simulation approach. Specifically, we sampled random starting positions for an initial set of cells at *t* = 0, then simulated their future locations using a defined tissue growth pattern. We further defined a spatial correlation pattern for the transcription rates *ρ*. After simulating the rate parameters, we calculated *u* and *s* using the analytical ODE solution. We added noise and simulated sparsity to match the properties of the real spatial transcriptomic data. We started with two simple types of tissue growth patterns, layered growth and radial growth. These mimic two different types of biologically plausible patterns–the brain cortex grows layer by layer, while other structures grow from the inside to the outside. Next, we used these patterns as building blocks and simulated two datasets (Fig. 2a). Note that all 2D streamplot visualizations in this paper, including Fig. 2a, represent “cell velocity” in physical coordinates, not abstract coordinates computed from UMAP as in previous RNA velocity papers. See the appendix for complete simulation details.

**Fig. 2.**
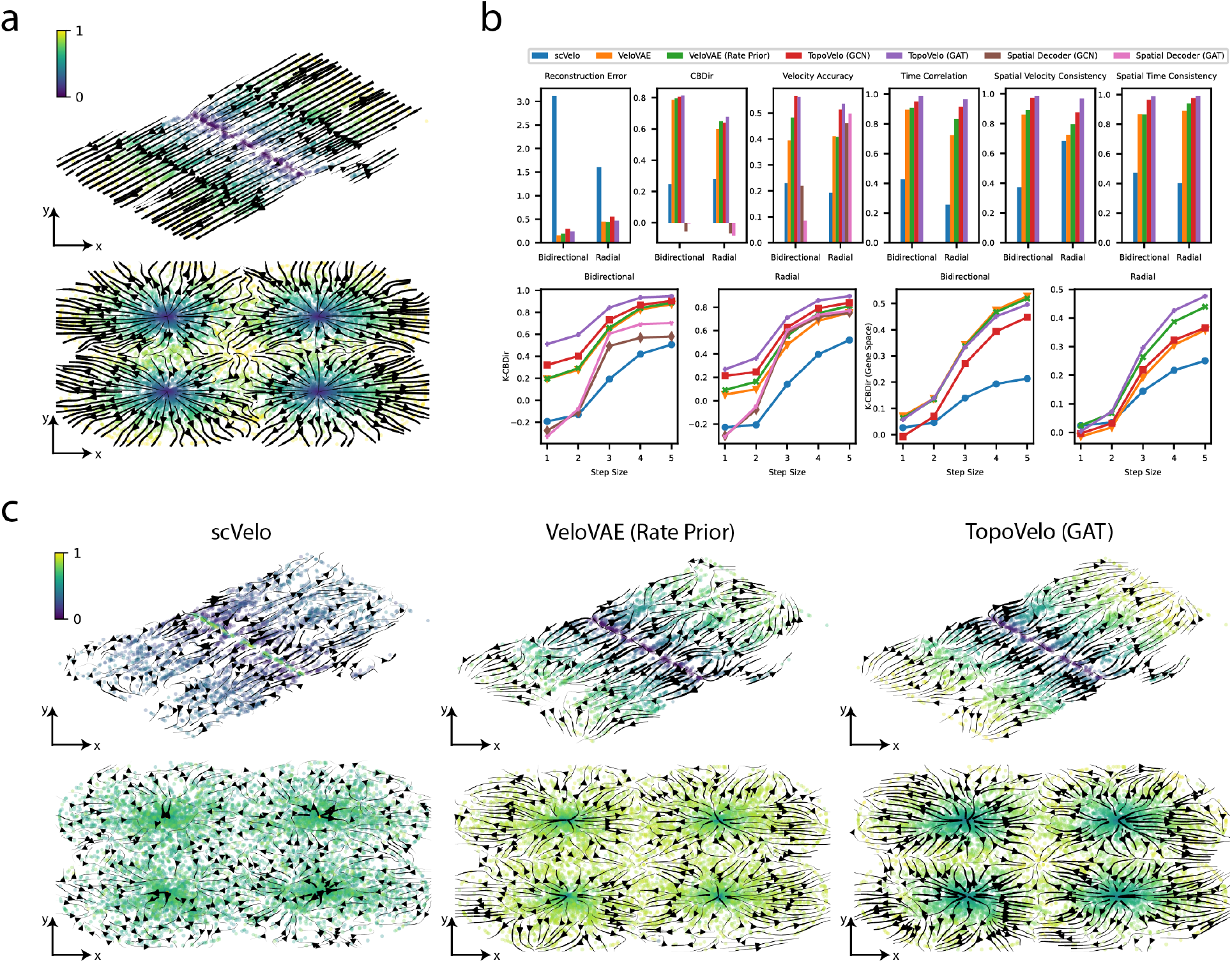
TopoVelo accurately models spatial dynamics in simulated datasets. **(a)** Simulated datasets with bidirectional(top) and radial(bottom) growth patterns. The true velocity stream is plotted over spatial coordinates colored by the true time values. Note that these stream plots indicate cell velocity in physical coordinates, not RNA velocity in abstract UMAP coordinates. **(b)** Quantitative performance. The top row shows six metrics used for benchmarking. Higher values are better for all metrics except for reconstruction error. The bottom row contains line plots showing *k*-CBDir using *k*-step neighbors with *k* up to five. Note that we computed two versions of *k*-CBDIR, one using spatial nearest neighbors and one using nearest neighbors in gene expression space. **(c)** Comparison of inferred cell velocity. Coordinates are colored by inferred time values.

To quantitatively evaluate our method, we computed several metrics and compared our performance with those from previous RNA velocity methods. We compared with scVelo[25], a widely used RNA velocity approach that fits each gene separately and assumes a single lineage, and VeloVAE[27], our previous method that fits a single shared latent time value for each cell and can model branching trajectories. Evaluation metrics include data reconstruction, cross-boundary direction correctness (CBDir)[26], velocity accuracy, time correlation, spatial velocity consistency and spatial time consistency. Data reconstruction error, represented by mean-squared error (MSE), quantifies how well a model fits the data. For evaluating cell velocities, we first employed cross-boundary direction correctness (CBDir)[26], a metric that measures the coherence of velocity flow between progenitor and descendant cell types. For computing CBDIR from simulated data, we divided cells into 5 clusters based on their true developmental time. We also introduced a new metric called *k*-CBDir by extending CBDir to include not only immediate neighbors but also *k*-step neighbors with *k* up to five. Additionally, we computed the average cosine similarity between inferred spatial velocity and the true cell velocity, which we refer to as “velocity accuracy”. Moreover, we calculated spatial velocity consistency–the average correlation of RNA velocity in a spatial neighborhood. To quantify spatial time consistency, we computed the local Moran’s I[39] of the time values inferred for nearby cells. These two spatial consistency metrics assess whether cell time and velocity vary smoothly among spatially adjacent cells instead of fluctuating rapidly over space.

Our results show that TopoVelo achieves significantly better performance in terms of CBDir, velocity accuracy, time correlation, spatial velocity consistency, and spatial time consistency (Fig. 2b). This indicates that incorporating spatial coordinates can substantially improve the estimation of cell times and velocities. The model with graph attention layers (GAT) consistently outperforms the model with spectral convolution layers (GCN). The reconstruction error is slightly higher than VeloVAE, but significantly better than scVelo. The cell velocity inferred by reconstructing spatial coordinates performs similarly to the velocity projected onto spatial coordinates for the radial growth pattern, but surprisingly performs worse on the bidirectional growth pattern. We suspect that this may be an artifact of the perfect symmetry of the simulated bidirectional growth pattern, which makes it harder for the spatial decoder to invert the function that maps from time to space. However, such perfect symmetry is unlikely to arise in real data, and indeed our results in the next section show that the spatial decoder performs well when applied to real datasets.

We next visualized cell velocity using velocity stream plots on spatial coordinates colored by inferred latent cell time (Fig. 2c). Since previous RNA velocity methods do not consider spatial information, we project the high dimensional RNA velocity to spatial coordinates following the same procedure as the scVelo method[25], except for replacing a gene-expression KNN graph with a spatial KNN graph. For TopoVelo, we visualize cell velocity from RNA velocity projection and spatial decoder separately. Among all three methods compared, TopoVelo has the strongest and most consistent velocity flow. The time inference from TopoVelo is also the closest to the ground truth, while the time from VeloVAE is mostly accurate yet less smooth spatially. These visual observations further support our previous quantitative results. Overall, our simulation study highlights the advantage of incorporating spatial context in modeling spatial transcriptomic data.

### 3.2 TopoVelo Accurately Infers Directions of Cell Migration and Differentiation in Developing Mouse Brain, Gut, and Lung

We next applied TopoVelo to real spatial transcriptomic datasets generated with the Slide-Seq v2[10] and Stereo-Seq[11] technologies. Slide-Seq measures transcriptome-wide mRNA expression levels at 10 *μ*m resolution, while Stereo-Seq measures transcriptome-wide mRNA expression at sub-*μ*m resolution. These technologies are ideal initial candidates for spatial velocity analysis because they have cellular resolution and capture both spliced and unspliced transcripts. We further chose a mouse cerebral cortex Slide-seq dataset from embryonic day 15 (E15) and mouse gut and lung Stereo-seq datasets from E13.5.

These real datasets exhibit spatial developmental patterns consistent with those designed in our simulation study. During the development of the mouse cerebral cortex, neurons integrate into cortical layers in a birthdate-dependent manner and follow a radial axis starting in the ventricular zone (VZ) and progressing through the subventricular zone (SVZ), intermediate zone (IZ), and cortical plate (CP)[40]. Embryonic mouse lung and gut form 3D tubular structures during development. During early embryonic development (E9.5 to E14.5), the developing gut and lung initially form pseudostratified epithelial tissue surrounded by splanchnic mesenchyme. From E9.5 to E14.5, the cells of the epithelium and mesenchyme rapidly proliferate, resulting in gut elongation and increase in the luminal area and girth of each tube[41,42].

We downloaded the authors’ Slide-Seq *u* and *s* counts and ran the published SAW pipeline to obtain *u* and *s* from Stereo-Seq data. In the absence of cell segmentation information, we binned the Stereo-Seq data at 20 *μ*m resolution. Additionally, the Stereo-Seq data contained cluster annotations only at the level of tissues (e.g., gut and lung), so we sub-clustered gut and lung spots using Leiden community detection and manually annotated clusters as epithelial or mesenchymal using previously published marker genes. We followed the same preprocessing steps used by previous RNA velocity methods: selecting highly variable genes, normalizing by library size, and smoothing *u* and *s* among nearest neighbors[25]. We tried smoothing by nearest neighbors calculated from spatial coordinates instead of principal components, but found this made results worse on both simulated and real data. We further tried fitting the model with no smoothing, but results were significantly worse in this case.

Although we lack detailed ground truth labels in the real data, we know the general spatial trend of tissue growth, and we know that inferred time and velocity should be spatially smooth. Additionally, we know that the cortex forms from the VZ to the upper layers (right to left in the Slide-Seq section) and that gut and lung tubes develop from the inside out. Thus, for the cortex dataset we used the x coordinate as a rough approximation of the expected temporal order. For the gut and lung datasets, we roughly approximated the expected temporal order by identifying the tissue border using Delaunay triangulation, then calculating the distance of each cell from the nearest border cell. We thus used this prior knowledge to quantitatively assess the performance of TopoVelo on these datasets by calculating CBDIR, *k*-CBDIR, time correlation, spatial velocity consistency, and spatial time consistency.

As shown in Fig.3a, TopoVelo with graph attention achieves the highest spatial time consistency, spatial velocity consistency, time correlation, and accuracy of velocity inference. We see a particularly significant advantage in terms of the spatial time consistency, most notably in the lung dataset, where both scVelo and VeloVAE have near-zero consistency. As with the simulated data, the data reconstruction error of TopoVelo is much better than scVelo but comparable to and slightly worse than that of VeloVAE. We also observed decent performance from the spatial decoder network (colored in pink), which learns a mapping from time and cell state to spatial coordinates. Our results clearly demonstrate TopoVelo’s outstanding performance in velocity inference from real data, especially when considering multi-step neighbors on the spatial graph.

We also qualitatively evaluated the results by visualizing velocity and latent time on tissue coordinates (Fig.3b, c). In the cerebral cortex, both TopoVelo and VeloVAE correctly captured the cell migration trend from the VZ zone to CP in the stream plots. However, TopoVelo produced more accurate cell time in the more differentiated layers including L5/6 and CP (Fig. 3b, c, first row). In contrast, scVelo not only showed a reversed migration direction in L5/6, CP and CR cells in the streamline plots, but also misidentified CR cells as the cell type of origin. In both streamline and latent time plots from the gut and lung data, TopoVelo exhibited a clear spatial pattern of outward radial growth. Although the developmental relation between mesenchymal and epithelial cells in early mouse gut and lung is not completely understood, previous literature concludes that from E9.5 to E14.5, pseudostratified intestinal epithelium and surrounding splanchnic mesenchyme rapidly proliferate, resulting in tube elongation and increase in both luminal area and girth[41,42]. This is consistent with our spatial velocity results, which predict that the tubes composed of epithelial cells (seen in cross-section in our 2D datasets) and surrounded by mesenchymal cells are growing outward. To further support this hypothesis, we plotted cell cycle scores[25] and found that cell cycle activity shows an inside-to-outside pattern (Fig. 3d).

**Fig. 3.**
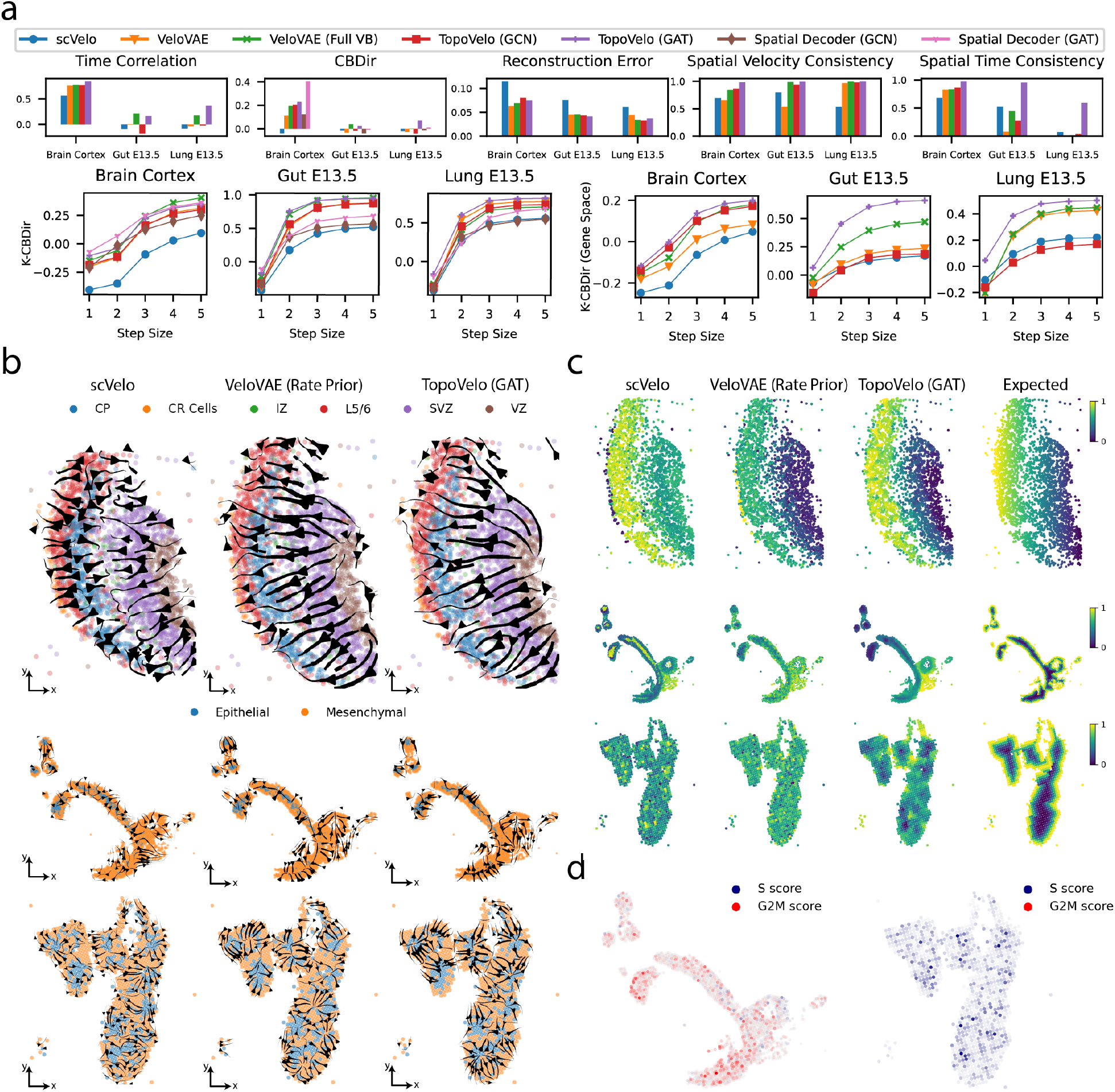
TopoVelo reveals spatial dynamics in mouse embryo development. **(a)** Performance comparison in terms of data reconstruction and velocity metrics. Higher values are better for all metrics except reconstruction error. Note that we computed two versions of *k*-CBDIR, one using spatial nearest neighbors and one using nearest neighbors in gene expression space. **(b)** Velocity stream plots shown on spatial (physical) coordinates. **(c)** Cell coordinates colored by inferred latent cell time. The last column entitled “Expected” shows a rough approximation of the expected time based on the knowledge that cortex grows from ventricular zone to upper layers and gut and lung form from the inside out. **(d)** G2/M phase and S phase scores computed by scVelo. A diverging color map, with G2/M phase in red and S phase in green, is used show cell cycle phase scores.

### 3.3 TopoVelo Enables Principled Modeling Across Multiple Slices and Time Points

A number of recent spatial transcriptomic protocols including Slide-seq and Stereo-seq have generated multi-slice data from samples at different developmental stages. Such multi-slice datasets can include sections from the same time and spatial plane but different tissue samples; slices from the same time and tissue sample but different spatial planes; or sections from different tissue samples across successive times. Our probabilistic framework can be readily extended to model such complex experimental designs in a principled fashion. As an example of how TopoVelo can model multi-slice data, we analyzed mouse gut Stereo-seq data from mice at E10.5, E11.5, E12.5 and E13.5. We know the embryonic day of each section, and we know that gut develops significantly during each of these embryonic days. Thus, we incorporated this information as a statistical prior on the time of each spatial gene expression profile. Encouragingly, we found that incorporating multiple slices and time labels improved the accuracy of the velocity inference. We compared the E13.5 gut results from only a single section with the results from jointly modeling the four sections spanning E10.5-E13.5 and using the time labels as a prior. When comparing the single-slice and multi-slice analyses, we used the same set of genes and calculated metrics only within a single slice (E13.5) for a fair comparison. The multi-slice analysis indeed improved the CBDIR, *k*-CBDIR, time correlation, and spatial time consistency metrics compared to the single-slice analysis (Fig. 4a). The multi-slice analysis also shows slightly lower but comparable spatial time consistency and slightly better reconstruction error. Qualitatively, our results show that TopoVelo infers velocity flow consistent with inside-to-outside development across all four time points (Fig. 4b). Finally, we demonstrate that by training a spatial decoder post hoc, we can obtain an explicit parametric function that predicts meaningful cell velocity (Fig. 4c). The spatial decoder produces a consistent outflow from the inner part of gut at all four time points, although it may oversmooth some local structures.

**Fig. 4.**
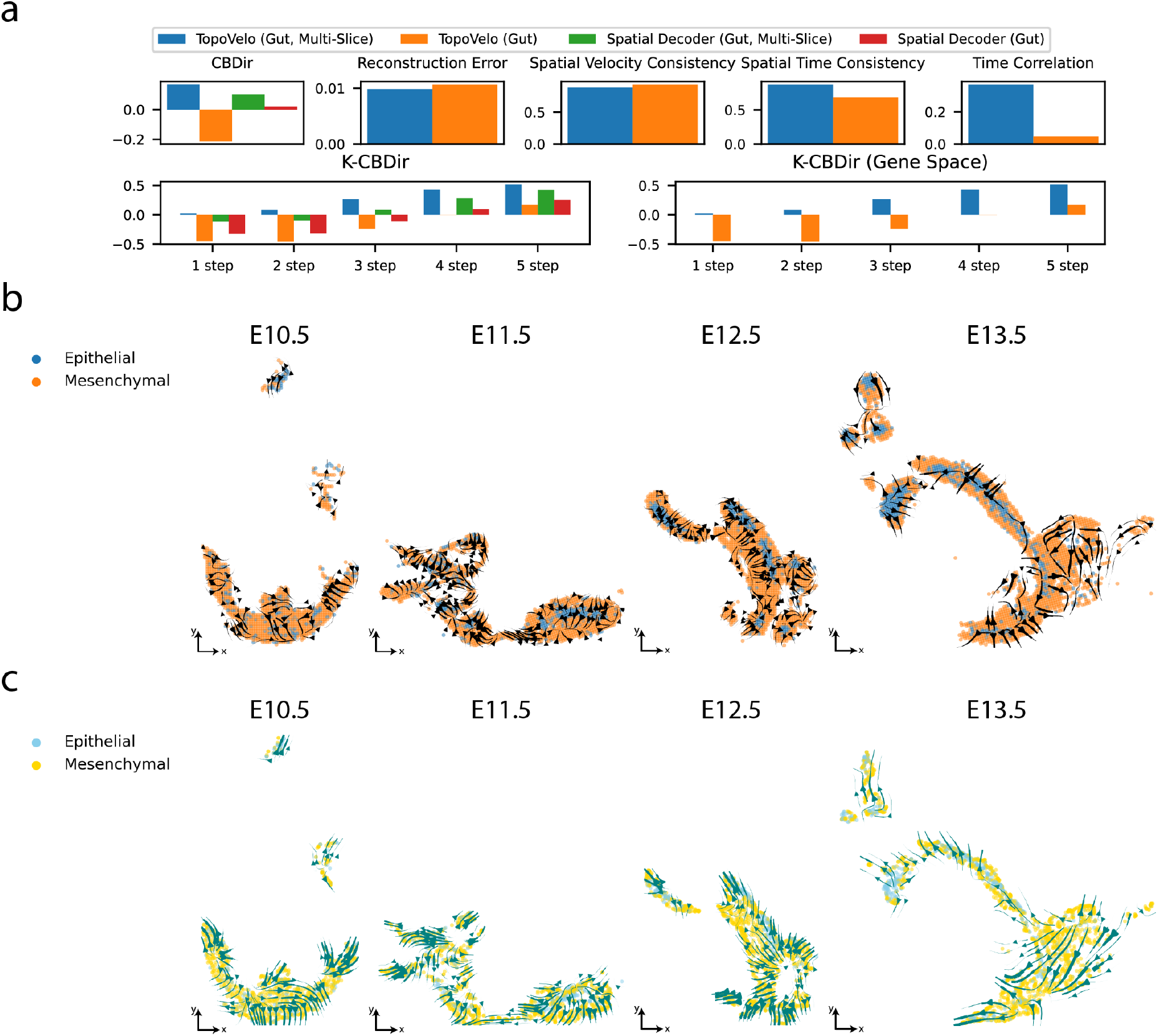
Multi-slice analysis incorporating time labels improves velocity inference. **(a)** Quatitative performance of two TopoVelo (GAT) models trained on a multi-slice and a single-slice dataset, respectively. Higher values indicate better performance for all metrics except for the reconstruction error. We used the expected temporal order from the single-slice data (E13.5) as the reference for computing time correlation. **(b)** Stream plot of mouse gut at each time point from TopoVelo. We apply the stream plot method from scVelo with the KNN graph replaced by a spatial graph. **(c)** Stream plot of mouse gut at each time point from the spatial decoder. The stream lines are plotted over predicted spatial locations from the spatial decoder.

## 4 Discussion

In this work, we consider the problem of modeling gene expression dynamics across an entire population of cells from spatial transcriptomic data. To our knowledge, TopoVelo is the first model that can use spatial transcriptomic data to link space and time in this way. We believe that this paper is only the first step in this exciting direction and there are a number of questions for future work to address.

First, our spatial decoder lacks an explicit mechanism describing cell migration. A potential improvement could be to incorporate domain knowledge about tissue growth and mechanistic models describing spatial movement of cells. Second, there is much room for improving multi-slice data integration and analysis. TopoVelo is capable of learning cell velocity from multi-slice data, but does not consider spatial coupling between two slices. With information from different stages, we envision predicting the entire process of tissue growth in terms of both RNA and cell velocity. Finally, we are excited to apply TopoVelo to gain a more comprehensive understanding of cell differentiation across a wide range of tissue environments.

## 5 Acknowledgements

This work was supported by the University of Michigan Rackham Predoctoral Fellowship to J.L. We thank Matthew Karikomi and Chao Gao for helpful discussions.

## 6 Code Availability

The source code is available at https://github.com/g-yichen/TopoVelo_RECOMB.

## References

1. Lane, S. W., Williams, D. A. & Watt, F. M. Modulating the stem cell niche for tissue regeneration. Nature biotechnology 32, 795–803 (2014).

2. Rock, J. R. et al. Basal cells as stem cells of the mouse trachea and human airway epithelium. Proceedings of the National Academy of Sciences 106, 12771–12775 (2009).

3. Angevine, J. & Sidman, R. L. Autoradiographic study of cell migration during histogenesis of cerebral cortex in the mouse. Nature 192, 766–768 (1961).

4. Berry, M. & Rogers, A. The migration of neuroblasts in the developing cerebral cortex. Journal of anatomy 99, 691 (1965).

5. Rakic, P. Mode of cell migration to the superficial layers of fetal monkey neocortex. Journal of Comparative Neurology 145, 61–83 (1972).

6. Pleasure, S. J. et al. Cell migration from the ganglionic eminences is required for the development of hippocampal gabaergic interneurons. Neuron 28, 727–740 (2000).

7. Wonders, C. P. & Anderson, S. A. The origin and specification of cortical interneurons. Nature Reviews Neuroscience 7, 687–696 (2006).

8. Butt, S. J. et al. The temporal and spatial origins of cortical interneurons predict their physiological subtype. Neuron 48, 591–604 (2005).

9. Bouffi, C. et al. In vivo development of immune tissue in human intestinal organoids transplanted into humanized mice. Nature biotechnology 1–8 (2023).

10. Rodriques, S. G. et al. Slide-seq: A scalable technology for measuring genome-wide expression at high spatial resolution. Science 363, 1463–1467 (2019).

11. Chen, A. et al. Spatiotemporal transcriptomic atlas of mouse organogenesis using dna nanoball-patterned arrays. Cell 185, 1777–1792.e21 (2022).

12. Cho, C.-S. et al. Microscopic examination of spatial transcriptome using seq-scope. Cell 184, 3559–3572.e22 (2021).

13. Russell, A. J. C. et al. Slide-tags: scalable, single-nucleus barcoding for multi-modal spatial genomics. bioRxiv (2023).

14. Ståhl, P. L. et al. Visualization and analysis of gene expression in tissue sections by spatial transcriptomics. Science 353, 78–82 (2016).

15. Thrane, K., Eriksson, H., Maaskola, J., Hansson, J. & Lundeberg, J. Spatially resolved transcriptomics enables dissection of genetic heterogeneity in stage iii cutaneous malignant melanoma. Cancer Research 78, 5970–5979 (2018).

16. Moncada, R. et al. Integrating microarray-based spatial transcriptomics and single-cell rna-seq reveals tissue architecture in pancreatic ductal adenocarcinomas. Nature Biotechnology 38, 333–342 (2020).

17. Chen, K. H., Boettiger, A. N., Moffitt, J. R., Wang, S. & Zhuang, X. Spatially resolved, highly multiplexed rna profiling in single cells. Science 348, aaa6090 (2015).

18. Wang, X. et al. Three-dimensional intact-tissue sequencing of single-cell transcriptional states. Science 361, eaat5691 (2018).

19. Janesick, A. et al. High resolution mapping of the breast cancer tumor microenvironment using integrated single cell, spatial and in situ analysis of ffpe tissue. bioRxiv 2022–10 (2022).

20. Welch, J. D., Hartemink, A. J. & Prins, J. F. Slicer: inferring branched, nonlinear cellular trajectories from single cell rna-seq data. Genome biology 17, 1–15 (2016).

21. Haghverdi, L., Büttner, M., Wolf, F. A., Buettner, F. & Theis, F. J. Diffusion pseudotime robustly reconstructs lineage branching. Nature methods 13, 845–848 (2016).

22. Qiu, X. et al. Reversed graph embedding resolves complex single-cell trajectories. Nature methods 14, 979–982 (2017).

23. Schiebinger, G. et al. Optimal-transport analysis of single-cell gene expression identifies developmental trajectories in reprogramming. Cell 176, 928–943.e22 (2019).

24. Klein, D. et al. Mapping cells through time and space with moscot (2023).

25. Bergen, V., Lange, M., Peidli, S., Wolf, F. A. & Theis, F. J. Generalizing rna velocity to transient cell states through dynamical modeling. Nature biotechnology 38, 1408–1414 (2020).

26. Gao, M., Qiao, C. & Huang, Y. Unitvelo: temporally unified rna velocity reinforces single-cell trajectory inference. Nature Communications 13, 6586 (2022).

27. Gu, Y., Blaauw, D. T. & Welch, J. Variational mixtures of ODEs for inferring cellular gene expression dynamics. In Chaudhuri, K.et al. (eds.) Proceedings of the 39th International Conference on Machine Learning, vol. 162 of Proceedings of Machine Learning Research, 7887–7901 (PMLR, 2022).

28. Cui, H., Maan, H., Taylor, M. D. & Wang, B. Deepvelo: Deep learning extends rna velocity to multi-lineage systems with cell-specific kinetics. bioRxiv (2022).

29. Qin, Q., Bingham, E., La Manno, G., Langenau, D. M. & Pinello, L. Pyro-velocity: probabilistic rna velocity inference from single-cell data. bioRxiv (2022).

30. Gayoso, A. et al. Deep generative modeling of transcriptional dynamics for rna velocity analysis in single cells. Nature Methods (2023).

31. La Manno, G. et al. Rna velocity of single cells. Nature 560, 494–498 (2018).

32. Li, C., Virgilio, M., Collins, K. L. & Welch, J. D. Multi-omic single-cell velocity models epigenome–transcriptome interactions and improves cell fate prediction. Nature Biotechnology 41, 387–398 (2023).

33. Kingma, D. P. & Welling, M. Auto-encoding variational bayes. In Bengio, Y. & LeCun, Y. (eds.) 2nd International Conference on Learning Representations, ICLR 2014, Banff, AB, Canada, April 14-16, 2014, Conference Track Proceedings (2014).

34. Kipf, T. N. & Welling, M. Variational graph auto-encoders (2016).

35. Märtens, K. & Yau, C. Basisvae: translation-invariant feature-level clustering with variational autoencoders. In Chiappa, S. & Calandra, R. (eds.) Proceedings of the Twenty Third International Conference on Artificial Intelligence and Statistics, vol. 108 of Proceedings of Machine Learning Research, 2928–2937 (PMLR, 2020).

36. Kipf, T. N. & Welling, M. Semi-supervised classification with graph convolutional networks. In International Conference on Learning Representations (2017).

37. Veličković, P. et al. Graph attention networks. In International Conference on Learning Representations (2018).

38. Kingma, D. P. & Ba, J. Adam: a method for stochastic optimization. In Bengio, Y. & LeCun, Y. (eds.) 3rd International Conference on Learning Representations, ICLR 2015, San Diego, CA, USA, May 7-9, 2015, Conference Track Proceedings (2015).

39. Anselin, L. Local indicators of spatial association—lisa. Geographical analysis 27, 93–115 (1995).

40. Lodato, S. & Arlotta, P. Generating neuronal diversity in the mammalian cerebral cortex. Annual review of cell and developmental biology 31, 699–720 (2015).

41. Wells, J. M. & Spence, J. R. How to make an intestine. Development 141, 752–760 (2014).

42. Kolev, H. M. & Kaestner, K. H. Mammalian intestinal development and differentiation—the state of the art. Cellular and Molecular Gastroenterology and Hepatology 16, 809–821 (2023).

