## Supplementary material for "Mapping Cell Fate Transition in Space and Time": methods_supplementary

### Appendix for: Mapping Cell Fate Transition in Space and Time

Yichen Gu[1*] Jialin Liu[2*] Chen Li[2]
Joshua D. Welch[2,3†]
[1]Dept. of Electrical and Computer Engineering, University of Michigan, Ann Arbor, MI, USA
[2]Dept. of Computational Medicine and Bioinformatics, University of Michigan, Ann Arbor, MI, USA
[3]Dept. of Computer Science and Engineering, University of Michigan, Ann Arbor, MI, USA

#### 1 Neural Network Details

**Spectral Graph Convolution.** Graph convolution is a form of message passing. The hidden state of a node is a weighted sum of contributions from all its neighbors. The convolution operation can be written in the following spectral form [1]:

$$\boldsymbol{H} = \tilde{\boldsymbol{D}}^{-\frac{1}{2}} \tilde{\boldsymbol{A}} \tilde{\boldsymbol{D}}^{-\frac{1}{2}} \boldsymbol{O} \boldsymbol{W} \tag{1}$$

where $\boldsymbol{O}$ is a cell-by-feature matrix, $\tilde{\boldsymbol{A}}$ is an adjacency matrix with self-loops, $\tilde{\boldsymbol{D}}$ is a diagonal degree matrix of $\tilde{\boldsymbol{A}}$, and $\boldsymbol{W}$ is a weight matrix learned from the data.

**Graph Attention.** Graph attention (GAT) applies the attention mechanism[2] to nodes in a graph. The original GAT paper mentioned several benefits, including that GAT is scalable to different neighborhood sizes and can be generalized to different graph structures[3]. For each node, GAT first linearly projects input features to another space, then computes an attention score over its neighborhood and aggregates the hidden information from the neighborhood. It can be written in the following form[3]:

$$\boldsymbol{h_i} = \sum_{j \in nbr(i)} k_{ij} \boldsymbol{W} \boldsymbol{o_i} \quad \text{where } k_{ij} = \frac{\exp(\boldsymbol{a}^T [\boldsymbol{W} \boldsymbol{o_i} \| \boldsymbol{W} \boldsymbol{o_j}])}{\sum_{j' \in nbr(i)} \exp(\boldsymbol{a}^T [\boldsymbol{W} \boldsymbol{o_i} \| \boldsymbol{W} \boldsymbol{o_{j'}}])} \tag{2}$$

We can enable multi-head attention by using multiple projection matrices $\boldsymbol{W}$ and concatenating the outputs.

#### 2 Simulation of unspliced and spliced mRNA data using coupled spatiotemporal dynamics.

In the simulation study, we extended the method from scVelo to include spatial information and couple it with temporal dynamics. We defined two different types of spatial dynamics in this

*Equal contribution

study, 1D bidirectional and 2D radial growth. The former mimics the process of a single layer of progenitor cells growing to its both sides, while the later is a case of cell expanding outwards to form concentric circles. These two cases are simple, but can be treated as building blocks for many real biological processes. For example, the VZ layer in the mouse brain cortex grows mostly in a constant direction to form subsequent layers. As for radial growth, epithelial cells in the inner part of the gut migrate outwards to become mesenchymal cells. Although these two simulated spatial dynamics differ, they share the same steps in our simulation. Therefore, we will describe our algorithm in a general sense, but refer the details to Table 1. In the following description, we always use $\mathcal{N}(\cdot,\cdot)$ to denote a uni- or multi-variate Gaussian distribution with mean and covariance matrix inside the parenthesis and $\mathcal{U}(a,b)$ to denote a uniform distribution in the interval $[a,b]$.

| | 1D Bidirectional | Radial | Notes |
|---|---|---|---|
| Initial Location | $x(0) \sim \mathcal{N}(0,\sigma_0^2)$<br>$y(0) \sim \mathcal{U}(-\frac{d}{2},\frac{d}{2})$ | $r(0) \sim \mathcal{U}(0,r_0)$<br>$\theta(0) \sim \mathcal{U}(0,2\pi)$ | |
| Spatial Dynamics | $v_x = v_0 + a_x t$<br>$v_y \sim B \cdot \mathcal{U}(v_{y,min}, v_{y,max})$ | $v_r = v_0 + a_r t$<br>$v_\theta \sim B \cdot \mathcal{U}(v_{\theta,min}, v_{\theta,max})$ | $B \sim \text{Beta}(\alpha,\beta)$ |
| Transcription Rate ($\rho$) | $\frac{1}{2}e^{\left(-\frac{(x-x^*)^2}{\sigma_x^2}-\frac{(y-y^*)^2}{\sigma_y^2}\right)}$<br>$+\frac{1}{2}e^{\left(-\frac{(x+x^*)^2}{\sigma_x^2}-\frac{(y+y^*)^2}{\sigma_y^2}\right)}$ | $e^{\left(-\frac{(r-r^*)^2}{\sigma_x^2}-\frac{(1-\cos(\theta-\theta^*))^2}{\sigma_\theta^2}\right)}$ | $(x^*,y^*) \sim \mathcal{U}([0,w] \times [-\frac{d}{2},\frac{d}{2}])$<br>$(r^*,\theta^*) \sim \mathcal{U}([0,R] \times [0,2\pi])$ |

Table 1: **Parametric model for spatial dynamics.** The table lists different forms of spatial velocity and spatially-dependent transcription rates used for simulating 1D bidirectional and 2D radial cases.

The simulation algorithm is composed of three parts, simulating the spatial dynamics, generating spatially dependent transcription rates, and simulating the temporal dynamics. In the first step, we can specify the initial number of cells at $t = 0$ and start with randomly sampling their initial positions. We assume cell development has a tree structure, i.e., a progenitor cell gives rise to one or multiple descendant cells. Each initial cell is the root of a tree, and we recursively build the tree. Given a progenitor cell in a tree with time $t$, it has $k$ children with time $t+\Delta t$, where $k$ is geometrically distributed and $\Delta t$ is exponentially distributed. This step stops when the number of cells in all the trees exceeds a threshold set by the user. Once cell time is sampled using the tree, we can predict their locations using the spatial dynamics model in Table1. After generating the spatial locations, the algorithm samples a spatially dependent transcription rate to resemble real spatial coordination of cells regarding gene expression. We assume the transcription rate has a peak at certain location and use a superposition of two heat kernels in the 1D bidirectional case. Similarly, we assume the transcription rate has a peak at a certain angle when the tissue is expanding radially and apply a heat kernel on the polar coordinates. Finally, we use the analytical form of the ODE solution (Eq. 3,4) to generate unspliced and spliced counts. To model technical noise, we added a Gaussian noise to the noiseless data and randomly set a proportion of entries to be zero as a result of dropout. We set the sparsity of our data to be 75%.

$$u(t) = u_0 \exp(-\beta\tau) + \frac{\rho\alpha}{\beta}(1 - \exp(-\beta\tau)) \tag{3}$$

$$s(t) = s_0 \exp(-\gamma\tau) + \frac{\rho\alpha}{\gamma}(1 - \exp(-\gamma\tau)) + \frac{\rho\alpha - \beta u_0}{\gamma - \beta}(\exp(-\gamma\tau) - \exp(-\beta\tau)) \tag{4}$$

### 3 Initialization of Gene-wise Latent Variable $\boldsymbol{w}$

In TopoVelo, $\boldsymbol{w}$ contains a set of categorical random variables representing if a gene is induced or repressed. Instead of using some uninformative Dirichlet prior, we can actually initialize these variables using the dynamical model.

We use the proportion of cells in the induction phase as an indicator of the gene being induced. First, we compute a gene-by-gene weighted adjacency matrix $\mathbf{A}$ where $A_{ij} = \frac{1}{2}[corr(\boldsymbol{U}_{:,i}, \boldsymbol{U}_{:,j}) + corr(\boldsymbol{S}_{:,i}, \boldsymbol{S}_{:,j})] + 1$, where $corr(\cdot,\cdot)$ denotes Pearson correlation coefficient and $\boldsymbol{U}_{:,i}, \boldsymbol{S}_{:,i}$ denotes unspliced and spliced counts of the $i$-th gene across all cells. We apply spectral clustering on genes using $\mathbf{A}$. The number of clusters is determined by using an empirical threshold on the singular values of $\mathbf{A}$ with a Gaussian noise assumption [4]. Let $\tilde{w}_g$ be the proportion of cells in the induction phase and $y_g \in \{1, 2, \ldots, K_y\}$ be the clustering result of the $g$-th gene. For each cluster $y$, we perform two Kolmogorov-Smirnov tests between $\mathcal{W}_y = \{w_g : y_g = y\}$ and Dirichlet$(5.0, 5.0)$. The two null hypotheses are

1. $\mathcal{W}_y$ is sampled from Dirichlet$(\psi_1, \psi_2)$ whose CDF is less than CDF of Dirichlet$(5.0, 5.0)$
2. $\mathcal{W}_y$ is sampled from Dirichlet$(\psi_1, \psi_2)$ whose CDF is greater than CDF of Dirichlet$(5.0, 5.0)$

If the first null hypothesis is not rejected, genes in cluster $y$ are likely to be inductive. We set $\psi_1 > \psi_2$ and reinitialize $\mathcal{W}_y$ by sampling from Dirichlet$(\psi_1, \psi_2)$. If the second null hypothesis is not rejected, genes in the cluster $y$ are likely to be repressive and we perform the same sampling but with $\psi_1 < \psi_2$. If both null hypotheses are rejected, we sample $\mathcal{W}_y$ from a mixture of two Dirichlet distributions mentioned in the previous two cases.

### References


[1] Kipf, T. N. & Welling, M. Semi-supervised classification with graph convolutional networks. In *International Conference on Learning Representations* (2017).

[2] Vaswani, A. *et al.* Attention is all you need. In Guyon, I. *et al.* (eds.) *Advances in Neural Information Processing Systems*, vol. 30 (Curran Associates, Inc., 2017).

[3] Veličković, P. *et al.* Graph attention networks. In *International Conference on Learning Representations* (2018).

[4] Gavish, M. & Donoho, D. L. The optimal hard threshold for singular values is $4/\sqrt{3}$. *IEEE Transactions on Information Theory* **60**, 5040–5053 (2014).
